# Genome analysis of *Lactobacillus plantarum* and *Lactobacillus brevis* isolated from traditionally fermented Ethiopian kocho and their probiotic properties

**DOI:** 10.1101/2025.09.04.674358

**Authors:** Guesh Mulaw, Teklemichael Tesfay, Tesfaye Sisay, Diriba Muleta, Débora Brito Goulart, Abdulafiz Musa, Nirupama Narayanan

## Abstract

Probiotics are essential for promoting health, with lactic acid bacteria (LAB) from traditional fermented foods like Ethiopian kocho offering valuable benefits. The objective of this study was to systematically analyse the genomic characteristics, bacteriocin production, and probiotic potential of LAB strains isolated from fermented Ethiopian kocho. The research involved isolating LAB from kocho, assessing their tolerance to acid and bile salts, evaluating antimicrobial activity, determining antibiotic susceptibility, and conducting whole-genome sequencing (WGS) to investigate genetic relatedness. Out of 150 LAB isolates, 7 (4.67%) exhibited remarkable acid tolerance, surviving at rates between 50.52–74.05% and 33.33–62.40% after 3 and 6 hours of exposure to pH 2, respectively. These seven acid-tolerant isolates also demonstrated exceptional resistance to 0.3% bile salt, maintaining survival rates ranging from 88.96% to 98.10% over 24 hours. In addition, the isolates displayed inhibitory effects against several important foodborne pathogenic bacteria, underscoring their potential as natural antimicrobial agents. Antibiotic susceptibility testing revealed that all isolates were susceptible to ampicillin, tetracycline, and erythromycin, whereas the most potent isolates exhibited significant resistance to kanamycin. Notably, four of the seven isolates showed resistance to streptomycin, while the remaining three were sensitive. The WGS analysis revealed that the isolates belonged to the *Lactobacillus* genus, including six *Lactobacillus plantarum* strains and one *Lactobacillus brevis* strain. Genomic analysis using the Bayesian Analysis of Gene Essentiality (BAGEL) tool predicted the presence of two class II bacteriocins across all seven strains, further supporting their potential as functional probiotic candidates. Overall, our findings highlight the probiotic potential of the seven *Lactobacillus* strains, demonstrating their acid and bile salt tolerance, antimicrobial properties, and genetic predisposition for bacteriocin production.

## 1. Introduction

Traditional fermented foods are integral to the human diet worldwide, with Africa being home to the most diverse array of such foods (1). These foods are geographically specific and have been developed through indigenous fermentation practices using locally sourced raw materials, both plant-based and animal-based (2, 3). Traditional fermentation processes are applied to various raw materials throughout the African continent (4). The consumption of traditional fermented foods and beverages is becoming increasingly popular worldwide for its associated health benefits, including but not limited to better cardiovascular health (5), improved lactose digestion (6), enhanced mental health (7), and strengthened immune system (8), among others. Additionally, fermented foods have been shown to support gut microbiota balance (9), reduce inflammation (10), and improve metabolic health (11), making them an essential part of a balanced diet. As a result, significant *in vivo* and *in vitro* research is being done worldwide to evaluate the effects of these foods on human health (12). Traditional fermented foods, with their rich cultural and geographical diversity, play a crucial role in promoting human health, especially in Africa where they are a key dietary component.

Probiotic microorganisms in fermented foods may contribute to host health by providing essential nutrients, supporting microbial growth, synthesizing enzymes, inhibiting pathogens, and modulating immune responses (13, 14). Probiotics are live microorganisms that confer health benefits to the host when administered in adequate quantities (15). The most prevalent probiotic lactic acid bacteria (LAB) include various species of *Lactobacillus* (e.g., *L. acidophilus*, *L. johnsonii*, *L. casei*, *L. rhamnosus*, *L. gasseri*, and *L. reuteri*) and the genus *Bifidobacterium* (e.g., *B. bifidum*, *B. animalis* subsp. *lactis*, *B. longum* subsp. *longum*, and *B. longum* subsp. *infantis*) (16). The probiotic efficacy of these microorganisms is often associated with host specificity, suggesting that strains isolated from the gut are more likely to be effective in probiotic applications (13). Furthermore, recent studies also highlight that the therapeutic potential of probiotics extends beyond gut health, influencing metabolic disorders, mental health via the gut-brain axis, and even modulating the gut microbiota diversity regarding beneficial bacteria (17, 18). Continued research into the mechanisms of action and specific strain-target interactions is essential to optimize their clinical and dietary applications.

Ethiopia is home to several fermented foods and beverages, typically produced through acid-alcohol fermentation processes (19). Previous studies have primarily focused on the microbiological aspects of fermentation involving milk and dairy products (20), as well as traditional staples such as injera (21). Kocho, a traditional Ethiopian food, is prepared from the decorticated and pounded pulp of the Ensete plant (*Ensete ventricosum*). The pulp is then mixed, kneaded into a mash, and fermented in a pit (22). Kocho, along with various fermented legume and vegetable products, and beverages, contributes to the diverse culinary landscape. The wide array of fermented foods and drinks consumed across different ethnic groups underscores notable cultural and dietary diversity. These fermented products, derived from both plant and animal sources, undergo biochemical and nutritional transformation through the activity of bacteria, yeast, and mold (19). Typically prepared at the household level, these products support local diets and nutritional intake, reflecting a deep-rooted culinary practice.

Genome analysis of *Lactobacillus plantarum* and *Lactobacillus brevis* isolated from traditionally fermented Ethiopian kocho has raised some controversies surrounding the accuracy of functional predictions and the genetic stability of probiotic strains. While genomic sequencing has allowed for the identification of promising probiotic traits, such as acid tolerance, bile salt resistance, and antimicrobial activity (23–25), the interpretation of these traits based solely on genomic data remains contentious. Researchers argue that *in silico* predictions may overestimate the probiotic potential, as they often fail to account for the variability in gene expression and functionality under real physiological conditions (26). Additionally, concerns have been raised regarding the genetic stability of *Lactobacillus* during prolonged storage and fermentation (27, 28), highlighting the need for more longitudinal studies to verify their long-term efficacy. Further experimental validation, including *in vivo* trials and long-term monitoring, is essential to fully assess the probiotic potential and stability of these strains in practical applications.

Although previous studies have explored the isolation, identification, and screening of LAB with antibacterial activity from traditional Ethiopian fermented foods, no comprehensive whole-genome sequencing (WGS) analysis has been conducted on probiotic strains isolated from foods such as kocho, particularly with a focus on bacteriocin-producing genes. This study is crucial as it fills the gap in existing research by providing a WGS analysis of probiotic strains isolated from kocho, offering insights into their genetic makeup, particularly bacteriocin-producing genes, which could unlock new avenues for developing natural antimicrobial agents and functional foods with enhanced health benefits. We hypothesized that LAB isolated from traditionally fermented Ethiopian kocho possess significant probiotic potential, including acid and bile salt tolerance, antimicrobial activity, and bacteriocin production, making them promising candidates for the development of functional foods. Our study aimed to investigate the probiotic potential of traditional Ethiopian fermented foods and their capacity to inhibit foodborne pathogens.

## 2. Materials and methods

### 2.1 Sample collection

A total of 15 kocho samples were collected from Welkite town and surrounding rural areas in Ethiopia, using a stratified random sampling approach to capture diversity in production practices and geographic origin. Samples were obtained from both household producers and local markets. Although the total sample size was limited to 15, stratified random sampling was employed to ensure diversity across production practices and geographic origins. This allowed for a representative cross-section of typical kocho products in the region. Approximately 200 g of each sample was aseptically collected into sterile containers. To maintain microbial integrity, the samples were transported to the Microbiology Laboratory at Addis Ababa University within 4 hours of collection, using insulated iceboxes equipped with temperature data loggers to ensure storage at 4 ± 1 °C. Upon arrival, they were stored at 4 °C and analysed within 48 hours to preserve microbial integrity.

### 2.2. Isolation of probiotic LAB from fermented kocho

To isolate LAB from the fermented Kocho samples, 10 g of each sample was aseptically homogenized in 90 mL of 0.1% (w/v) sterile peptone water using a stomacher for 2 minutes. Serial ten-fold dilutions were prepared up to 10⁻⁶. Aliquots (0.1 mL) from dilutions 10⁻³ to 10⁻⁶ were spread in duplicate onto De Man, Rogosa, and Sharpe (MRS) agar plates. The plates were incubated anaerobically at 37 °C for 24 hours in anaerobic jars (AnaeroPack-Anaero, Mitsubishi Gas Chemical Co.) with an oxygen indicator to ensure anaerobic conditions. After incubation, 3 to 5 colonies with distinct morphological characteristics (based on size, shape, edge, and pigmentation) were selected from each plate, streaked onto fresh MRS agar plates, and incubated under the same anaerobic conditions for an additional 24 hours to obtain pure isolates. Sterile controls were included to monitor for contamination throughout the process.

### 2.3 Probiotic characteristics of the isolated LAB

#### 2.3.1. Acid and bile salts

To assess acid tolerance, overnight LAB cultures were harvested by centrifugation at 5000 rpm for 10 min, washed twice with sterile phosphate-buffered saline (PBS), and resuspended in MRS broth adjusted to pH 2.0, 2.5, or 3.0 using 1N HCl. Suspensions were incubated at 37 °C for 3 and 6 hours under anaerobic conditions. For bile salt tolerance, isolates were inoculated into MRS broth containing 0.3% (w/v) bile salt and incubated at 37 °C for 24 hours. Survival rate (%) was calculated using the formula (29):

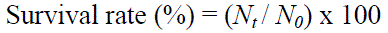

where *N_0_* and *N_t_* represent the viable biomass (measured by OD₆₀₀ or CFU/mL) at 0 and specified time points, respectively. All experiments were conducted in triplicate, and data were analysed using one-way analysis of variance (ANOVA) followed by Tukey’s test (p < 0.05).

#### 2.3.2. Antimicrobial activity

Antimicrobial activity of the LAB cell-free supernatants (CFS) was evaluated using the agar well diffusion method (30), with slight modifications. The test pathogens included *Staphylococcus aureus* ATCC 25923, *Escherichia coli* ATCC 25922, *Listeria monocytogenes* (clinical isolate), and *Salmonella enterica* serovar Typhimurium (clinical isolate), all obtained from the Ethiopian Public Health Institute (EPHI). LAB isolates were cultured in MRS broth at 37 °C for 24 hours, and cultures were centrifuged at 6000 rpm for 10 minutes at 4 °C. The supernatants were filtered through 0.22 µm filters to obtain sterile CFS. To minimize the effect of organic acids, the pH was adjusted to 6.5 using 1N NaOH. For some tests, catalase (1 mg/mL) was added to eliminate hydrogen peroxide activity. Pathogenic strains were grown in Brain Heart Infusion (BHI) broth for 18–24 hours and adjusted to 0.5 McFarland turbidity (∼10⁸ CFU/mL). 100 µL of each bacterial suspension was spread evenly on sterile nutrient agar (NA) plates (90 mm diameter). Once dried, wells (5 mm in diameter) were made using a sterile cork borer, and 100 µL of CFS was added to each well. Plates were incubated at 37 °C for 24 hours. Zones of inhibition around the wells were measured in millimetres. All experiments were performed in triplicate. Sterile MRS broth served as a negative control, and ampicillin (10 µg) discs were included as positive controls.

#### 2.3.3. Antibiotic susceptibility tests

The antibiotic susceptibility of LAB isolates (adjusted to ∼10⁸ CFU/mL) was assessed using the disc diffusion method (30), following modified Clinical and Laboratory Standards Institute (CLSI) (31) protocols adapted for LAB, and supported by the European Food Safety Authority (EFSA) guidance (32) and prior studies (33, 34). LAB cultures were spread onto MRS agar plates supplemented with 0.5% glucose. The following antibiotic discs were used: ampicillin (10 μg), erythromycin (15 μg), streptomycin (10 μg), kanamycin (25 μg), and tetracycline (30 μg) (Oxoid, UK). Plates were incubated anaerobically at 37 °C for 24 to 48 hours. Inhibition zones were measured using a digital caliper, and results were interpreted as resistant (R ≤ 15 mm), intermediate (I = 16 to 20 mm), or susceptible (S ≥ 21 mm), based on modified criteria adapted from CLSI (31). *E. coli* ATCC 25922 was used as a quality control strain. All tests were conducted in triplicate, and data were expressed as mean ± standard deviation (SD).

### 2.4. Whole-genome sequencing (WGS) of probiotic LAB isolates

#### 2.4.1. DNA extraction

Genomic DNA was extracted from fresh overnight cultures of LAB using a modified cetyltrimethylammonium bromide (CTAB) method (35). A 0.5 mL aliquot of each culture was mixed with 40% sterile glycerol in a 1.5 mL microcentrifuge tube and incubated at 37 °C with shaking at 230 rpm until the culture reached an optical density (OD₆₀₀) of approximately 1.0. Cells were harvested by centrifugation at 4000 rpm for 15 minutes, and the resulting pellet was resuspended in 10 mM Tris-HCl buffer (pH 8.5). The suspension was divided equally into three sterile 1.5 mL microcentrifuge tubes and centrifuged again at 4000 rpm for 2 minutes, after which the supernatant was discarded.

The pellets were washed with 400 µL of wash buffer (10 mM Tris-HCl, 50 mM NaCl, 1 mM ethylenediamine tetraacetic acid (EDTA), pH 8.0, and the supernatant was removed after centrifugation. Each pellet was resuspended in 400 µL of elution buffer (10 mM Tris-HCl, 0.1 mM EDTA, pH 8.5), followed by the addition of 20 µL lysozyme (100 mg/mL), 5 µL mutanolysin (10 U/µL), 45 µL proteinase K (20 mg/mL), and 1 µL RNase A (10 mg/mL). The mixture was incubated at 37 °C for 15–30 minutes for enzymatic digestion of the cell wall and nucleic acids.

After enzymatic treatment, 70 µL of 10% sodium dodecyl sulfate (SDS) was added, and the tubes were gently inverted to mix. The samples were incubated at 65 °C for 10 minutes. Following this, 100 µL of 5 M NaCl and 100 µL of pre-warmed 10% CTAB solution in 0.7 M NaCl were added. The tubes were vortexed gently and incubated again at 65 °C for 10 minutes.

For DNA extraction, 500 µL of chloroform:isoamyl alcohol (24:1) was added to each tube, vortexed for 10 seconds, and centrifuged at 12,000 × g for 5 minutes. The upper aqueous phase was carefully transferred to a new sterile microcentrifuge tube. To precipitate DNA, cold isopropanol was added at 0.6 × the volume of the aqueous phase, and samples were incubated at −20 °C for at least 30 minutes. DNA was then pelleted by centrifugation at 12,000 × g for 10 minutes, washed with 500 µL of cold 70% ethanol, air-dried for 5 minutes at room temperature, and resuspended in 50 µL of elution buffer.

To remove residual RNA, 1 µL of RNase A (10 mg/mL) was added to each tube, and samples were incubated at 37 °C for 30 minutes. Further purification was achieved by adding 5 µL of 3 M sodium acetate (pH 8.0) and 100 µL of cold 99% ethanol, followed by gentle mixing and centrifugation at 12,000 × g for 2 minutes. The supernatant was discarded, and the DNA pellet was washed with 70 µL of cold 70% ethanol, air-dried, and resuspended in 50 µL of elution buffer.

The integrity of extracted genomic DNA was verified by electrophoresis on a 1% agarose gel stained with ethidium bromide. DNA concentration was measured using the Qubit dsDNA Broad Range Assay Kit (Thermo Fisher Scientific, USA) according to the manufacturer’s protocol (36). DNA purity was evaluated using a NanoDrop spectrophotometer (Thermo Fisher Scientific, USA) by calculating the absorbance ratios at 260/280 nm and 260/230 nm.

#### 2.4.2. Whole genome sequencing

Following DNA extraction, WGS was performed at the Earlham Institute (Norwich, UK). Low-Input Transposase-Enabled (LITE) libraries were constructed according to the facility’s standard protocol and sequenced using the Illumina HiSeq 4000 platform, generating 2×150 bp paired-end reads. Raw reads underwent quality control and preprocessing using BBDuk (part of the BBTools suite) (37). Adapter sequences were trimmed from the 3′ ends, and bases with Phred quality scores below 3 were removed from both ends. Reads shorter than 100 bp or with an average Phred score below 20 were discarded. Quality of the trimmed reads was assessed using FastQC (38). The filtered reads were normalized to a coverage depth ranging from 2× to 100× to reduce computational load and potential assembly errors. *De novo* genome assembly was performed using SPAdes version 3.8.1 (39), which is optimized for small-genome microbial assemblies. Default k-mer lengths were used, and assembly quality was evaluated based on N50 values, contig length, and total assembly size. Assembled genome sequences were functionally annotated using the Rapid Annotations using Subsystems Technology (RAST) server (40), available at http://rast.nmpdr.org/ and http://bioinf.spbau.ru/en/spades. This pipeline assigns gene functions and categorizes genes into biological subsystems, providing high-quality genome-scale metabolic and functional annotations.

#### 2.4.3. Bacteriocin prediction

Putative bacteriocin gene clusters were predicted using the Bacterial Genome Annotation and Comparison Lab’s wysiwyg Genome Finder 4 (BAGEL4) pipeline (41), a web-based genome mining tool specialized for the identification of bacteriocin operons and associated biosynthetic genes. Assembled genome contigs were submitted to the BAGEL4 web server (http://bagel4.molgenrug.nl/databases.php), which utilizes a combination of Hidden Markov Models (HMMs), curated databases, and motif-based searches to identify and classify bacteriocin gene clusters, including Classes I (e.g., lantibiotics), II (e.g., small heat-stable peptides), and III (e.g., large heat-labile proteins).

In addition to BAGEL4’s automated pipeline, a local BLASTx search was performed against bacteriocin protein sequences obtained from the BAGEL4 reference database. Contigs were aligned using BLASTx with an E-value threshold of 1e-10 and a minimum sequence identity of 80%. Only contigs meeting these criteria were classified as encoding putative bacteriocin proteins. The combination of BAGEL4 predictions and high-confidence BLASTx alignments allowed robust identification and classification of bacteriocin-associated sequences within the genome.

### 2.5. Data analysis

All experiments were conducted in triplicate, and the results reported as mean ± SD. Statistical analyses were carried out using SAS software (version 9.2; SAS Institute Inc., Cary, NC, USA). Where appropriate, ANOVA was used to assess differences among groups, and p-values less than 0.05 were considered statistically significant. Specific statistical tests (e.g., one-way ANOVA, Tukey’s HSD post hoc test, or Student’s t-test) were selected based on data distribution and experimental design. For genomic analyses, *de novo* genome assembly was performed using the SPAdes genome assembler (version 3.8.1) (39). Annotation of assembled genomes was conducted using the RAST pipeline (40), which provides functional classification of genes based on curated subsystems.

## 3. Results

### 3.1. In vitro characterization of probiotic properties

#### 3.1.1. Resistance to low pH

The screening of 90 probiotic LAB isolates revealed varying degrees of acid tolerance. After a 3-hour exposure to different pH conditions, 7 isolates (7.78%) demonstrated viability at pH 2, while another 7 isolates (7.78%) survived exposure to pH 2.5. A higher number of isolates, 13 (14.44%), were found to be tolerant to a less acidic environment of pH 3. Extending the exposure time to 6 hours resulted in a decrease in the number of viable isolates at the higher pH levels. Specifically, all 7 isolates that were viable at pH 2 and pH 2.5 after 3 hours remained viable, but the number of isolates surviving at pH 3 decreased to 9. The overall analysis showed that 7 (7.78%) of the total 90 LAB isolates exhibited survival at all three pH values (2, 2.5, and 3) after both 3 and 6 hours of exposure (Figure 1).

**Figure 1.**
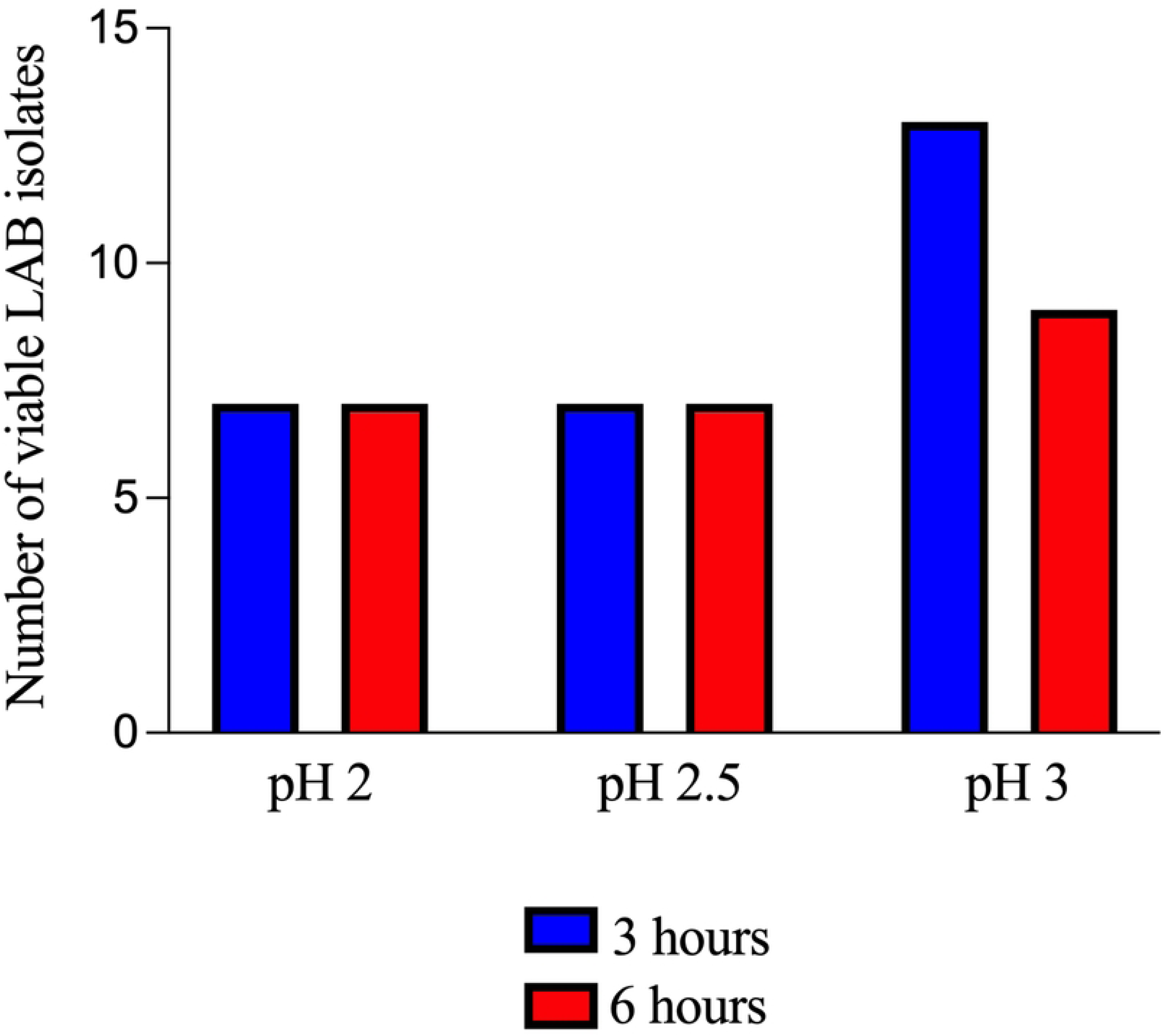
Acid tolerance patterns of LAB at different pH values after 3 and 6 hours of exposure.

Quantitative analysis of the survival rates revealed significant differences among the viable LAB isolates under the various acidic conditions (p < 0.05). The survival rates ranged from 33.33±1.03% to 97.35±1.40%. After 3 hours of incubation at pH 2, isolate K135 demonstrated the highest survival rate at 74.05±1.59%. This was followed by K082 (69.92±1.56%) and K071 (64.82±2.01%). In contrast, K070 exhibited the lowest tolerance, with a survival rate of 50.52±0.69%. Similarly, at pH 2 for 6 hours, K135 remained the most resilient, maintaining a survival rate of 62.40±1.22%. Three isolates showed survival rates below 50%, ranging from 33.33±1.03% to 45.80±1.26%.

At less acidic conditions (pH 2.5 and pH 3), the survival percentages were generally higher. Isolate K135 consistently showed high tolerance, with survival rates of 86.74±0.72% at pH 2.5 and 97.35±1.40% at pH 3 after 3 hours of exposure. Conversely, isolate K040 showed the lowest survival rates at these pH values: 70.16±0.73% at pH 2.5 and 83.69±0.73% at pH 3. Although most of the isolates survived, the survival percentages decreased with increasing acidity and prolonged exposure time. At pH 2.5 for 6 hours, survival rates ranged from 50.40±1.43% to 79.01±1.06%. The survival rates at pH 3 for 6 hours ranged from 63.99±0.70% to 90.31±1.59%. Among the seven isolates tested at pH 3 for 6 hours, K135 displayed the highest survival rate (90.31±1.59%), followed by K071 (87.44±2.30%) and K070 (82.70±0.72%). Table 1 shows the survival rates of probiotic LAB at various pH levels and 0.3% bile salt concentrations.

**Table 1.** Percentage survival of probiotic LAB at different pH levels and 0.3% bile salt.

| Isolates | pH tolerance |  |  |  |  |  | Bile tolerance |
| --- | --- | --- | --- | --- | --- | --- | --- |
|  | 3 h |  |  | 6 h |  |  | 24 h |
|  | pH 2 | pH 2.5 | pH 3 | pH 2 | pH 2.5 | pH 3 | 0.3% |
| K070 | $50.52 \pm 0.69$ | $74.91 \pm 1.03$ | $92.78 \pm 1.03$ | $33.33 \pm 1.03$ | $50.40 \pm 1.43$ | $82.70 \pm 0.72$ | $95.72 \pm 1.19$ |
| K040 | $55.48 \pm 1.4$ | $70.16 \pm 0.73$ | $83.69 \pm 0.73$ | $45.80 \pm 1.26$ | $53.97 \pm 1.07$ | $63.99 \pm 0.70$ | $88.96 \pm 1.18$ |
| K135 | $74.05 \pm 1.59$ | $86.74 \pm 0.72$ | $97.35 \pm 1.40$ | $62.40 \pm 1.22$ | $79.01 \pm 1.06$ | $90.31 \pm 1.59$ | $98.10 \pm 1.18$ |
| K122 | $54.26 \pm 0.78$ | $67.05 \pm 1.55$ | $88.24 \pm 1.18$ | $45.22 \pm 1.36$ | $60.85 \pm 1.78$ | $76.49 \pm 1.57$ | $89.72 \pm 1.33$ |
| K071 | $64.82 \pm 2.01$ | $77.72 \pm 1.76$ | $91.46 \pm 1.81$ | $51.42 \pm 3.07$ | $69.01 \pm 1.26$ | $87.44 \pm 2.30$ | $94.74 \pm 0.48$ |
| K072 | $62.42 \pm 1.66$ | $76.42 \pm 1.42$ | $89.94 \pm 0.98$ | $51.26 \pm 2.13$ | $64.94 \pm 1.19$ | $77.83 \pm 1.89$ | $92.34 \pm 1.62$ |
| K082 | $69.92 \pm 1.56$ | $79.56 \pm 1.19$ | $92.58 \pm 0.7$ | $51.43 \pm 1.37$ | $67.06 \pm 1.19$ | $79.30 \pm 0.78$ | $92.58 \pm 1.17$ |

#### 3.1.2. Tolerance to bile salts

The acid tolerance of probiotic LAB isolates was highly dependent on both the environmental pH and the duration of exposure. As illustrated in Fig. 1, there was a significant decrease in the number of viable isolates at lower pH values and with prolonged exposure times. The quantitative data presented in Table 1 further supports this observation, with average survival rates decreasing from over 90% at pH 3.0 to as low as 33.33% at pH 2.0. Notably, isolate K135 exhibited consistently superior acid tolerance across all conditions, achieving the highest survival rates. This high level of resilience makes K135 a promising candidate for further investigation as a potential probiotic. The findings underscore the importance of rigorous, strain-specific tolerance testing to select the most effective probiotic candidates.

#### 3.1.3. Evaluating antimicrobial activity

The antimicrobial activity of the crude extract from each LAB isolate against common foodborne pathogens was assessed based on inhibition zone diameters (Table 2). The average inhibition zones ranged from 15 to 20 mm, indicating significant antimicrobial potential. Notably, isolate K070 exhibited the most pronounced antibacterial activity against *S. aureus* ATCC 25923, *L. monocytogenes*, *E. coli* ATCC 25922, and *S. typhimurium*, with inhibition zones ranging from 17.33 to 20 mm in diameter. In contrast, isolate K082 exhibited the lowest inhibition zones, ranging from 15 to 17.33 mm. Among the seven potential probiotic LAB candidates, isolate K122 produced the smallest inhibition zone (15.33 mm) against *L. monocytogenes*, while K070 demonstrated the largest inhibition zone (20 mm) against the same pathogen. Furthermore, K070 and UK072 exhibited minimal inhibition zones of 16 mm and 16.33 mm, respectively, while isolate K040 produced the largest inhibition zone (20 mm) against *S. aureus* ATCC 25923. Isolate K082 presented the smallest inhibition zone (15 mm) against *E. coli* ATCC 25922 and *S. typhimurium*, whereas K070 displayed the greatest inhibition zone (20 mm) against *E. coli* ATCC 25922. Likewise, isolate K135 demonstrated the largest inhibition zone (18.67 mm) against *S. typhimurium* (Table 2).

**Table 2.** Antimicrobial activities of LAB isolates against common foodborne pathogens.

| Number | Isolate | Diameter of inhibition zone (mm) |  |  |  |
| --- | --- | --- | --- | --- | --- |
|  |  | *L. monocytogenes* | *S. aureus* | *E. coli* | *S. typhimurium* |
| 1 | K070 | 20.00±1.00 | 16.00±1.00 | 20.00±1.00 | 17.33±0.58 |
| 2 | K040 | 17.33±1.53 | 20.00±1.00 | 19.67±0.58 | 17.67±1.53 |
| 3 | K135 | 17.33±1.53 | 18.00±1.00 | 18.67±0.58 | 18.00±1.00 |
| 4 | K122 | 15.33±0.58 | 18.33±0.58 | 17.00±1.00 | 16.00±1.00 |
| 5 | K071 | 16.33±1.53 | 17.67±1.53 | 17.67±1.53 | 16.67±0.58 |
| 6 | K072 | 19.00±1.00 | 16.33±1.53 | 17.67±0.58 | 16.33±1.53 |
| 7 | K082 | 17.33±1.53 | 17.00±1.00 | 15.00±1.00 | 15.00±1.00 |

### 3.2. Antibiotic susceptibility testing

Seven LAB isolates were evaluated for their susceptibility or resistance to various antibiotics. The selected panel of probiotic LAB isolates were sensitive to commonly used antibiotics, including tetracycline, ampicillin, and erythromycin. However, all seven isolates exhibited resistance to kanamycin. Further, three isolates were sensitive to streptomycin, while the remaining four showed resistance to this antibiotic (Table 3).

**Table 3.**
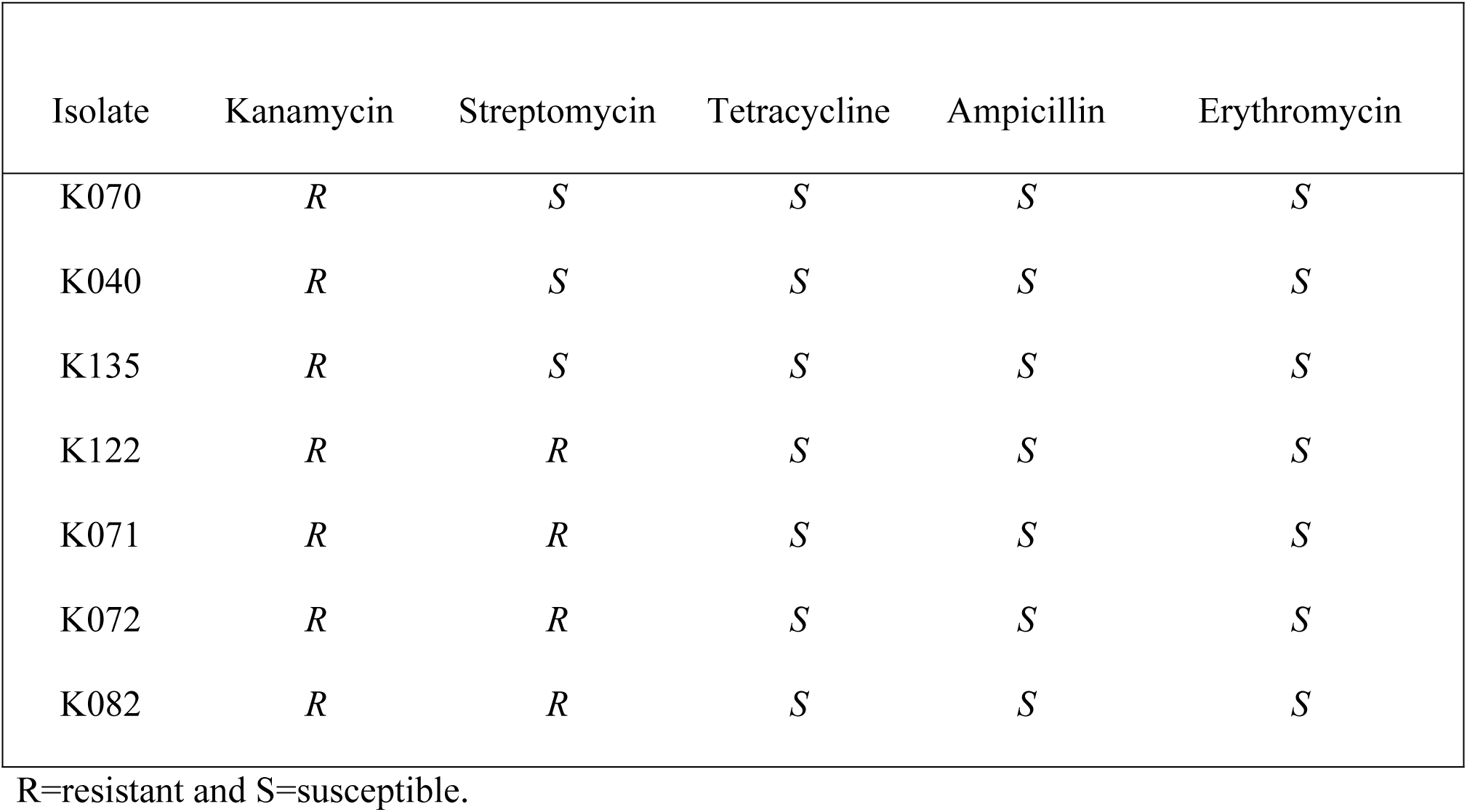
Antibiotic susceptibility profile of potential probiotic LAB isolates.

| Isolate | Kanamycin | Streptomycin | Tetracycline | Ampicillin | Erythromycin |
| --- | --- | --- | --- | --- | --- |
| K070 | *R* | *S* | *S* | *S* | *S* |
| K040 | *R* | *S* | *S* | *S* | *S* |
| K135 | *R* | *S* | *S* | *S* | *S* |
| K122 | *R* | *R* | *S* | *S* | *S* |
| K071 | *R* | *R* | *S* | *S* | *S* |
| K072 | *R* | *R* | *S* | *S* | *S* |
| K082 | *R* | *R* | *S* | *S* | *S* |

R=resistant and S=susceptible.

### 3.3. Identification of probiotic LAB isolates by WGS

The seven probiotic LAB isolates were further characterized through WGS (Table 4). Analysis of the 16S rRNA and *recA* gene sequences revealed that these isolates exhibited 99 to 100% sequence homology with known bacterial species, specifically *Lactobacillus plantarum* and *Lactobacillus brevis.* Notably, isolate K072 displayed 100% similarity to *L. plantarum* DK0 22T (Table 4). Among the seven strains, six were identified as *L. plantarum*, with guanine and cytosine (GC) content ranging from 44.4 to 45.9%. The other isolate, *L. brevis*, had 45.8% GC content. Among the *L. plantarum* strains, ATCC 14917 exhibited the highest GC content (45.9%), while *L. plantarum* ATCC 14917 and *L. plantarum* DK0 22T had the lowest GC percentage (44.4%).

**Table 4.** Genomic features of *Lactobacillus* species used in genomic comparisons.

| Isolates | Species | Strains | Identities (%) | Size (bp) | GC (%) | Contigs | CDS |
| --- | --- | --- | --- | --- | --- | --- | --- |
| K070 | *L. plantarum* | ATCC 14917 | 99.87 | 2,987,637 | 44.7 | 113 | 3,054 |
| K040 | *L. plantarum* | ATCC 14917 | 99.83 | 3,425,501 | 44.4 | 56 | 3,428 |
| K135 | *L. plantarum* | ATCC 14917 | 99.91 | 3,221,109 | 44.5 | 46 | 3,199 |
| K122 | *L. plantarum* | ATCC 14917 | 99.74 | 3,089,460 | 44.5 | 183 | 3,144 |
| K071 | *L. brevis* | ATCC 14869 | 99.54 | 2,460,308 | 45.8 | 103 | 2,547 |
| K072 | *L. plantarum* | DK0 22T | 100.00 | 3,426,101 | 44.4 | 115 | 3,429 |
| K082 | *L. plantarum* | ATCC 14917 | 99.74 | 2,481,105 | 45.9 | 45 | 2,476 |

Genome sequences of the six *L. plantarum* strains ranged from 2,481,105 to 3,426,101 bp, with 2,476 to 3,429 predicted coding sequences (CDSs). The genome sequence of the *L. brevis* strain was 2,460,308 bp in length and had 2,547 candidate CDSs. Notably, the *L. plantarum* DK0 22T (K072) genome was the largest, spanning 3,426,101 bp with 3,429 candidate CDSs, while *L. brevis* ATCC 14869 (K071) had the smallest genome, containing 2,460,308 bp and 2,547 CDSs. Overall, the genomes of the seven probiotic strains consisted of 45 to 183 contigs, with *L. plantarum* ATCC 14917 (K082) having the lowest number of 45 and *L. plantarum* ATCC 14917 (K122) displaying the highest number of 183 contigs.

### 3.4. Bacteriocin gene cluster identification

Among the seven potential probiotic strains, two harboured genes encoding components necessary for bacteriocin biosynthesis (Table 5). Specifically, *Lactobacillus plantarum* ATCC 14917 strains K040 and K135 were predicted to produce putative class II bacteriocins. Overall, the BAGEL tool identified one class II bacteriocin in each of these two genomes. Further, all blast alignments were manually inspected to confirm that the alignments were consistent with the presence of bacteriocin proteins.

**Table 5.** Potential probiotic strains sharing different classes of bacteriocin proteins.

| Isolate code | Species | Strains | Bacteriocin |  |  |
| --- | --- | --- | --- | --- | --- |
|  |  |  | Class I | Class II | Class III |
| AAUK040 | *Lactobacillus plantarum* | ATCC 14917 | - | + | - |
| AAUK135 | *Lactobacillus plantarum* | ATCC 14917 | - | + | - |

## 4. Discussion

Fermented foods are increasingly recognized as reservoirs of beneficial microbes with probiotic potential, particularly LAB, which contribute to food safety, preservation, and host health (42, 43). Among these, kocho, a traditional Ethiopian fermented food derived from *Ensete ventricosum*, remains largely unexplored for its microbial diversity and probiotic potential. The indigenous fermentation practices involved in its production provide a unique ecological niche for the evolution and enrichment of functionally relevant microorganisms (44). Previous studies have emphasized the importance of identifying novel, robust LAB strains from traditional fermented foods, especially those adapted to stress conditions such as low pH and bile salts, and capable of producing antimicrobial compounds (45, 46). In the present study, LAB strains were isolated from Kocho samples collected from different regions around Welkite, Ethiopia. The isolates were screened for probiotic properties including acid and bile tolerance, antimicrobial activity, antibiotic susceptibility, and the presence of bacteriocin biosynthesis genes. WGS was used to confirm strain identity and functional potential. Seven LAB strains, identified as *Lactobacillus plantarum* and *Lactobacillus brevis*, exhibited strong probiotic traits including acid and bile tolerance, as well as antimicrobial activity against key pathogens. Among them, isolate K135 stood out for its exceptional resilience and the presence of a class II bacteriocin gene cluster. This highlights the potential of kocho as a valuable source of indigenous probiotic strains that may be suitable for future applications in functional foods or as natural biopreservatives.

In this study, only seven out of 90 LAB isolates (7.78%) demonstrated notable acid tolerance, surviving exposure to pH 2 for up to 6 hours, with survival rates ranging from 33.33% to 97.35%. This wide range of tolerance suggests strong strain-dependent variability in acid resilience, which is critical for probiotic survival through the gastric environment (47). The ability of these isolates to endure such acidic conditions implies their potential to remain viable in the human stomach, an essential criterion for effective probiotic function. These results align with a previous study on *Lactobacillus* species from traditionally fermented Ethiopian foods, where strains showed survival rates ranging from 38.40% to 73.29% after 6 hours at pH 2 (48). Similarly, another study reported survival rates between 52% and 112% among 25 *Lactobacillus* strains after 4 hours of exposure to pH 2 (49). However, our results contrast with those of Li and collaborators, who reported that LAB strains from Chinese sourdough presented survival rates below 50% after 3 hours in simulated gastric juice at pH 2, with some strains losing viability (50). Conversely, another investigation found that most *Lactobacillus* spp. had survival rates exceeding 90% after 3 hours at pH 2, which was higher than the average found in the current study (51). These discrepancies may stem from differences in strain origin, adaptation to specific fermentation environments, or methodological variations in acid stress testing. The identification of acid-tolerant strains, particularly isolate K135, highlights the potential of Ethiopian fermented foods as valuable sources of robust probiotic candidates and supports the need for region-specific screening to uncover unique microbial resources.

The seven LAB isolates demonstrated considerable tolerance to acidic conditions, with survival rates ranging from 67.05% to 97.35% at pH 2.5 and from 50.40% to 90.31% at pH 3, over incubation periods of 3 and 6 hours. These results indicate a high level of acid resistance, which is a key prerequisite for probiotic functionality, as it reflects the ability of microorganisms to survive passage through the human stomach. Other studies also found elevated survival rates of *Lactobacillus* spp. after 3 hours of incubation. For example, *Lactobacillus* strains isolated from the traditional Korean fermented millet beverage Omegisool exceeded 88% survival rate at both pH 2.5 and 3 (52), while strains isolates from Iranian fermented dairy products survived at rates ranging from 71% to 76% at pH 2.5 (53). Similarly, resistance to acidic environments has also been observed in feces of piglets, in which all *Lactobacillus plantarum* strains demonstrated high tolerance at pH 3.0 for 3 hours (54). Notably, our findings confirm the high acid tolerance of certain isolates even after prolonged exposure (6 hours), which is more stringent than the 3-hour exposure commonly reported in the literature. This suggests that the LAB strains identified in this study may possess enhanced acid-adaptive mechanisms, possibly due to their origin in the naturally acidic environment of fermented kocho. Furthermore, the observed survival rates surpass those reported by Vinothkanna and Sekar (2020), where only 37% of isolates survived above 70% at pH 2.5, highlighting the robustness of the Ethiopian LAB strains (55). Given that food typically resides in the stomach for approximately 3 hours, where pH can fluctuate based on diet and physiological factors (56), the high survival of these isolates under such acidic conditions supports their practical application as orally administered probiotics.

In addition to acid tolerance, the seven LAB isolates from this study exhibited strong resistance to bile salts, further indicating their potential for survival in the small intestine. After 24 hours of exposure to 0.3% bile salts, all isolates maintained high viability, with survival rates ranging from 88.96% (K040) to 98.10% (K135). These results align with an earlier study reporting high bile salt tolerance among *Lactobacillus* strains, with survival rates ranging from 88% to 92% (53). Comparable resistance levels have been found in *Lactobacillus* isolated from traditional fermented foods such as Ethiopian shamita and kocho (57) and Jordanian fermented products (58), suggesting that traditional fermentation may contribute to the development of probiotic candidates. Bile salts are natural antimicrobial due to their detergent-like characteristics, which can disrupt bacterial cell membranes (59). Therefore, bile salt tolerance is widely considered a crucial functional advantage for probiotic selection. Resistance to bile salts found in LAB strains in the current study may be attributed to the activity of bile salt hydrolase (BSH), an enzyme responsible for deconjugating bile acids into less toxic forms (60). Although some *Lactobacillus* strains exhibited low tolerance (3% to 36%) to bile salts, most LAB strains isolated from fermented Ethiopian kocho demonstrated both acid and bile salt resistance, indicating their potential as probiotics capable of surviving the challenging human gastrointestinal environment.

Another important trait of functional probiotic strains is their antibacterial potential against common foodborne pathogens. In the current study, LAB isolates demonstrated varying levels of inhibition against *Staphylococcus aureus* ATCC 25923, *Listeria monocytogenes*, *Escherichia coli* ATCC 25922, and *S. typhimurium*. The antimicrobial activity was classified based on the diameter of inhibition zones, with a diameter less or equal to 9 mm indicating weak activity and a diameter higher or equal to 12 mm indicating strong inhibition. Among the isolates, K070 exhibited the strongest inhibition overall, with zones of 20.00 ± 1.00 mm against both *L. monocytogenes* and *E. coli*. Most isolates demonstrated moderate to strong inhibition across all pathogens, with inhibition zones ranging from 15.00 to 20.00 mm, highlighting their potential as antibacterial agents. These findings are in line with previous studies. Bassyouni and colleagues reported *Lactobacillus* strains from Egyptian dairy products exhibiting inhibition zones ranging from 17 to 21 mm against *E. coli* and *S. typhimurium* (61). Similarly, Choi and colleagues found that certain *Lactobacillus* strain completely inhibited several foodborne pathogens, including *E. coli* O157 ATCC 35150, *Salmonella enteritidis* KCCM 12021, *Salmonella typhimurium* KCTC 1925, and *Staphylococcus aureus* (62). Ryu and Chang demonstrated that *Lactobacillus plantarum* NO1, isolated from kimchi, inhibited *S. aureus* and *S. typhi* with inhibition zones ranging from 13.15 mm to 16 mm (63). Haghshenas and colleagues reported inhibition zones ranging from 11.7 mm to 13.7 mm against similar pathogens derived from *L. plantarum* strains in dairies (53). Overall, the LAB isolates in the current study exhibited notable antibacterial activity against major foodborne pathogens, underscoring their potential as effective probiotic candidates for pathogen control.

The antibiotic resistance pattern of LAB is an important factor in assessing their safety for probiotic use. In this study, all LAB isolates showed intrinsic resistance to kanamycin. Likewise, previous studies reported that *Lactobacillus* spp. are naturally resistant to aminoglycosides (64), which is explained by the impermeability of their cell membrane and the lack of electron transport chain required for drug absorption (64, 65). On the other hand, all LAB isolates were susceptible to tetracycline, ampicillin, and erythromycin, aligning with the typical susceptibility profile of LAB reported in previous studies (64, 66, 67). However, the study found variability related to streptomycin susceptibility: isolates K070, K040, and K135 were susceptible, while K122, K071, K072, and K082 were resistant. This heterogeneity is commonly found among *Lactobacillus* strains and may be related to strain-specific differences or environmental adaptations (68, 69). Resistance to aminoglycosides and vancomycin in LAB is generally intrinsic and does not present a risk of horizontal gene transfer (HGT) to commensal microbiota in the human gut (70). These findings corroborate existing literature on antibiotic susceptibility patterns for these *Lactobacillus* species, which support their use in fermented foods without posing significant public health risks (71). Generally, the intrinsic resistance patterns of LAB to certain antibiotics, together with their lack of risk for HGT, support their safety for use in fermented foods.

WGS confirmed that all seven probiotic candidates belonged to the genus *Lactobacillus*, comprising six *L. plantarum* ATCC 14917/DK022T strains and one *L. brevis* ATCC 14869 strain. Genome sizes of these isolates ranged from approximately 2.42 to 3.53 Mb and a GC content from 44.4% to 45.9%. Likewise, genomes of *L. paracasei* strains have been reported to have a GC content of 46.3% and a genome size of 3 Mb (72). Another study identified several *Lactococcus* species with variable genome sizes and GC content, with *L. garvieae* strains having the highest GC content of approximately 38.80% and the smallest genome sizes ranging from1.95 Mb to 2.60 Mb (73). A comparative study done by Kelleher and colleagues involving 30 lactococcal strains reported an average genome length of 2.428 Mb, ranging from 2.250 to 2.589 Mb (74). The same reseachers also found that *L. lactis* genome harbored an average of 2,344 predicted CDS, ranging from 1,947 to 2,643 CDS (74). Genomic sequence analysis of *L. plantarum* strains E2C2 and E2C5 done by another group revealed lengths of 3,603,563 bp and 3,615,168 bp, respectively (75). These genomes had GC contents of 43.99% and 43.97% and contained 3,289 and 3,293 candidate CDS (75). Despite the different isolation sources, the high genomic similarity of these isolates suggests their selective adaptation to the gut environment.

Bacteriocins are potent antimicrobial peptides produced by specific microorganisms, including LAB, that exhibit activity against closely related organisms. Production of bacteriocins is strain and culture-condition dependent (76). Among the seven genomic strains analysed for the prediction of putative bacteriocins, BAGEL4 identified two class II bacteriocins. In a related study, BAGEL4 predicted one bacteriocin for each of the three classes in *Lactococcus lactis* NCDO 2118 (73). The same study detected genes encoding components involved in bacteriocin synthesis, regulation, and related hypothetical proteins in the genome of *Lactobacillus rhamnosus* L156.4 (73). While some LAB strains are recognized as probiotic, others may be potentially probiotic or simply fermentative cultures widely distributed in nature with potential applications in the food industry (77). Likewise, not all probiotic LAB strains produce bacteriocins, underscoring the need for comprehensive screening to identify strains capable of producing the desired bacteriocins.

## 5. Conclusion

Our study has successfully identified and characterized seven LAB strains isolated from traditionally fermented Ethiopian kocho as having strong probiotic potential. These isolates exhibited high levels of tolerance to acidic and bile environments, suggesting that they are potentially tolerant for survival in the human GI. In addition to being tolerant to acidic and bile conditions, these LAB showed broad-spectrum antimicrobial activity against common foodborne pathogens and were generally susceptible to different antibiotics (Table 3). WGS confirmed the identity of these isolates to be *Lactobacillus plantarum* and *Lactobacillus brevis* and indicated the presence of genes encoding class II bacteriocins, which further indicates potential natural antimicrobial properties. Based on our findings, the LAB strains used in this traditional fermentation process can be considered for inclusion in functional foods and probiotic products. Our study acknowledges certain limitations that should be considered when interpreting the results. While the *in vitro* probiotic potential of the isolates was well-established, the study lacks long-term clinical or animal trials to confirm the strains’ efficacy and safety *in vivo*. Moreover, the reliance on computational tools like BAGEL4 for predicting bacteriocin production without experimental verification poses a limitation, as such predictions may not accurately reflect the actual bacteriocin production capabilities of the strains in real-world conditions. Future studies could expand on this research by exploring the *in vivo* probiotic efficacy of the isolated *Lactobacillus* strains, particularly their impact on gut health and immunity in animal models or human clinical trials. Additionally, investigating the stability of these probiotics under various storage conditions and their potential for scaling up production in fermented foods could offer valuable insights. Further genomic analysis could focus on identifying novel antimicrobial peptides or enzymes produced by these strains.

## Acknowledgements

The authors acknowledge the Department of Microbial, Cellular, and Molecular Biology, Addis Ababa University and Quadram Institute Bioscience, Norwich Research Park, Norwich, United Kingdom for their provision of laboratory and facilitation of the work.

